# ENaC contributes to macrophage dysfunction in cystic fibrosis

**DOI:** 10.1101/2024.11.06.622340

**Authors:** John Moran, Courtney Pugh, Nevian Brown, Ashley Thomas, Shuzhong Zhang, Emily McCauley, Amelia Cephas, Chandra L. Shrestha, Santiago Partida-Sanchez, Shasha Bai, Emanuela Bruscia, Benjamin T. Kopp

**Author notes:** To whom correspondence should be addressed: Benjamin Kopp, Emory Children’s Center, 2015 Uppergate Drive, Atlanta, GA 30322. **Author contributions:** Experimental performance: JM, CP, NB, AT, SZ, EM, AC, CLS. Study design and analysis: JM, CP, SZ, CS, BTK. Manuscript writing and editing: JM, CP, NB, SZ, EM, AC, CLS, SPS, SB, EB, BTK. Statistical analysis: SB, BTK. Grant support: BTK, SPS, EB. Recruitment: BTK.

## Abstract

**Background:** Cystic fibrosis (CF) is a chronic systemic disease caused by dysfunctional or absent cystic fibrosis transmembrane conductance regulator (CFTR). CFTR is expressed in human immune cells and plays a role in regulating innate immunity both directly and indirectly. Besides CFTR, research indicates that the epithelial sodium channel (ENaC) also contributes to dysfunction in CF airway epithelial cells. However, the impact of non-CFTR ion channel dysfunction on CF immune responses is not yet fully understood. A precise understanding of how CF immune function is regulated by ion channels may allow antibiotic-and mutation-agnostic treatment approaches to chronic bacterial infection and inflammation. Therefore, we hypothesized that ENaC is aberrantly expressed in CF macrophages and directly contributes to impaired phagocytic and inflammatory functions.

**Methods:** ENaC expression was characterized in human immune cells isolated from CF and non-CF blood donors. Monocyte-derived macrophage (MDM) function and bacterial killing was tested in the setting of ENaC modulation.

**Results:** Baseline expression of ENaC in human CF MDMs, lymphocytes, and granulocytes was increased at both the transcript and protein level relative to non-CF controls and persisted after exposure to bacteria. Inhibition of CFTR in non-CF MDMs resulted in ENaC overexpression.

CFTR modulator treatment reduced but did not eliminate ENaC overexpression in CF MDMs. Interestingly, ENaC inhibition with Amiloride increased CFTR expression. Amiloride-treated CF MDMs also showed normalized ROS production, improved autophagy, and decreased pro-inflammatory cytokine production. Finally, results from an ion channel microarray indicated that sodium channel expression in CF MDMs normalized after Amiloride treatment with minimal effect on other ion channels.

**Discussion:** ENaC is overexpressed in CF immune cells and is associated with abnormal macrophage function. ENaC modulation in immune cells is a novel potential therapeutic target for infection control in CF, either in combination with CFTR modulators, or as a sole agent for patients not currently eligible for CFTR modulators.

## INTRODUCTION

Cystic fibrosis (CF) is a chronic systemic disease classically attributed to dysfunctional or absent cystic fibrosis transmembrane conductance regulator (CFTR), an ion channel that regulates chloride and bicarbonate transport. CFTR is expressed in human immune cells and can directly and indirectly regulate innate immunity.^1-5^ The role of other ion channel dysfunction in CF immune responses remains less clearly defined. In addition to CFTR, many studies demonstrate that ENaC contributes to CF airway epithelial cell dysfunction.^6-8^ In contrast, other studies suggest that ENaC activity is not altered in CF.^9,10^ Further, how restoration of CFTR through small molecule or other therapeutic approaches will impact ENaC remains unclear. ENaC is composed of three homologous subunits (SCNNα, β, and γ), with a 4th subunit (δ) identified in specific human tissues. Overexpression of the beta subunit (SCNN1β) recapitulates the mucoinflammatory aspects of human CF lung disease in murine models, demonstrating the potential importance of ENaC in disease progression.^11^ Various therapies to restore normal ENaC expression in the CF airway are being pursued.^6^ Despite these efforts, ENaC is poorly described in CF immune cells, presumably due to its association as an epithelial-based channel. Recently, ENaC was described in CF monocytes and shown to influence inflammatory signaling.^12^ Prior studies showed that ENaC is also present in non-CF lymphocytes^13^ and neutrophils.^14^

A precise understanding of how CF immune function is regulated by non-CFTR ion channels would allow antibiotic- and mutation-agnostic treatment approaches to chronic bacterial infection and inflammation. To this end, we hypothesized that ENaC is aberrantly expressed in CF macrophages and directly contributes to dysfunctional phagocytic and inflammatory functions. Our studies demonstrate that ENaC is over-expressed in a variety of CF cell types including monocyte-derived macrophages (MDMs). Further, inhibition of ENaC helps improve critical deficits in CF macrophage-mediated bacterial killing though restoration of intracellular killing pathways. These findings have broad implications for the role of sodium transport via ENaC in regulating macrophage host responses against bacteria in CF and other lung diseases.

## MATERIALS AND METHODS

### Human participants

CF and non-CF study participants were recruited as approved by the Nationwide Children’s Hospital Institutional Review Board (IRB16-01020) and Emory University IRB (STUDY00004965). Written informed consent and/or assent was received prior to participation. People with CF (pwCF) were included if at baseline health and no prior history of bone marrow or lung transplant. People without CF were included if they had no underlying chronic illnesses and were age- and gender-matched to PWCF.

### Reagents

Amiloride (Sigma) 1-100uM, capsazepine (Sigma) 30uM, SPX101 (R. Tarran) 10uM, EIPA (Sigma) 10uM, Elexacaftor/tezacaftor/ivacaftor (ETI, Vertex) 3µM for each component, anti-LC3B (Sigma, L8918-200), anti-SCNN1β (abcam, ab28668), anti-SCNN1Δ (abcam, ab196737), anti-SCNN1γ (abcam ab3468), Alexa Fluor 488 FITC dye (Thermo Fisher), DAPI (Sigma Aldrich).

### Macrophage isolation

Peripheral blood was obtained during routine labs in EDTA tubes with monocytes isolated per our prior methods using Ficoll gradients.^1,5^ Isolated monocytes were cultured in RPMI plus 20% human AB serum and differentiated for 5 days at 37°C into MDMs as previously described, without additional cytokine stimulation.^15,16^ MDM purity and integrity was confirmed by microscopy and flow cytometry. MDMs were plated in a monolayer culture with fresh RPMI and 10% AB serum, rested for 4 days, with subsequent downstream experimentation. In some experiments macrophages were exposed to CF airway supernatants (ASN) obtained from filtered human sputum.

### Ion channel expression by flow cytometry

Intracellular staining of ENaC and CFTR was performed as previously described.^5^ In brief, MDMs (1.0×10^6^/per condition) were incubated in fixation buffer (Invitrogen, 00822249) for 20 min at RT and then treated with permeabilization buffer (Invitrogen, 00833356) for 15 min after 2 washes with cold PBS. Cells were centrifuged at 390×g for 5 min at 4°C and resuspended in 500 μL blocking buffer containing 5% goat serum (gibco, 16210072) in permeabilization buffer for 20 min at RT. Before flow cytometry, 100uL permeabilization buffer with ENaC (SCNN-1β, abcam) or CFTR (CFTR-596 from CFTR Antibody Distribution Program) antibodies at a dilution of 1:100 was added for 30 min at RT in the dark, and then stained with 100 uL permeabilization buffer with Alexa Fluor488 Goat anti-mouse antibody at a dilution of 1:300 (Molecular Probes, A31619) for 20 min at RT.

### Immunoblotting

ENaC and LC3 expression by immunoblotting was performed as previously described for MDM samples.^1,5^ In brief 3x 10e^6^ MDMs were plated and infected with *B. cenocepacia* clinical isolates overnight (MOI 2). Protein was then harvested after lysis in buffer containing a protease inhibitor (Sigma, 1183617001 Roche). Cell lysates were immunoblotted for β-actin and SCNN1γ. Relative expression level of proteins was quantified by densitometry of immunoblots, using ImageJ software (National Institutes of Health).

### Confocal with quantitation

Confocal microscopy was performed as previously described.^5^ In brief, 1 ×10^6^ macrophages were cultured on 12 mm glass cover slips in 24-well plates and stained with DAPI and anti-CFTR/ENaC/LC3b. Cells were fixed and permeabilized with 0.1% saponin buffer post staining. Blinded fluorescence images were obtained using a Zeiss LSM 800 (Carl Zeiss Inc., Thornwood, NY). Colocalization analysis was determined by ImageJ software version (National Institutes of Health).

### qRT-PCR/ion channel array

RNA was harvested and purified from MDMs via a total RNA purification kit (Norgen Biotek #17200) and concentrated with a RNA clean and concentrator kit (Zymo research #R1013) following standard protocols as previously described.^5,17^ Reverse transcription reactions were performed using a high-capacity cDNA reverse transcription kit according to the manufacturer protocol (Thermo fisher #4368814). qPCR amplification was then performed using SCNN1β primers or a TaqMan Array 96-well plate Custom Format48 (Thermo Fisher Scientific # 4391526) containing 45 predefined probes & 3 endogenous controls.^5^

### ROS production

ROS production via a DCF assay was performed as previously described.^2,5^ In brief, MDMs were seeded into 96-well plates at a density of 0.8 ×10^6^ per well in a volume of 200 μl fresh RPMI plus 10% human AB serum. After resting for 4 days, cells were treated with ETI or ENaC inhibitor compounds. Cells were then incubated for 30 min in Dulbecco’s PBS with HEPES 10mM, human serum albumin 1mg/ml plus 0.1% glucose and 10% DCF was added for 30 min. PMA 200 μM or *B. cenocepacia* MOI 10 was used to stimulate MDMs and fluorescence intensity was measured every 2 min for 2 hours using a Synergy H1 fluorescence microplate reader (BioTek) at an excitation wavelength of 485 nm and a 515 nm emission wavelength.

### Autophagosome detection

Autophagy induction was measured via the CYTO-ID autophagy detection kit (Enzo) using the manufacturer’s protocol. 2 × 10e^6^ MDMs were plated in 12 well plates and specific wells treated with ETI in warm RPMI or RPMI alone for 3 days. On the third day, Amiloride 100 µM and SPX-101 10 µM were added to respective wells. Six hours after treatment, cells were infected with *Burkholderia cenocepacia* at an MOI of 2. Simultaneously, rapamycin (autophagy inducer) and chloroquine (lysosomal inhibitor) were added as positive controls. Cells were incubated overnight and processed for live cell analysis of % autophagosome formation by flow cytometry with a proprietary CYTO-ID solution that selectively labels accumulated autophagic vacuoles.

### Bacterial killing

A colony forming unit (CFU) assay was performed as previously described.^1,5^ Briefly, 0.8 ×10^6^ MDMs were cultured on 24-well plates and treated with ETI, amiloride, SPX-101, or combinations before adding CF clinical isolates of MRSA, *Pseudomonas. aeruginosa*, and *B. cenocepacia* for 2 hours at 37°C. Extracellular bacteria were killed by gentamicin (50 μg/ml) for 40 min at 37°C. MDMs were replaced with fresh media and incubated overnight. MDMs were lysed in PBS 0.1% Triton X-100 (Acros Organics) and bacteria load was quantified by plating serial dilutions on LB agar plates and analyzed for CFUs.

### Cytokine production

Cytokines were measured via a LEGENDplex™ Human Inflammation Panel 1 according to the manufacturer’s protocol (Biolegend).

### Statistical analysis

Statistical analyses were completed with GraphPad Prism software (version 10). For any two-group comparisons (e.g. non-CF vs CF), two sample unpaired t-tests were used for parametric normally distributed data or the Mann-Whitney U test for non-parametric normally distributed data were used during single comparisons. For an overall comparison with more than groups, one-way ANOVA with post-hoc Tukey correction test was used for multiple comparison adjustments as detailed in figure legends.

## RESULTS

### ENaC is over-expressed in CF immune cells

Due to ENaC’s proposed interactions with CFTR, we hypothesized that ENaC may be upregulated in CF immune cells akin to the airway epithelium. To test this hypothesis, we detected ENaC in primary human CF MDMs by western blot, flow cytometry, microscopy, and qRT-PCR. CF macrophages had significantly higher detection of ENaC (all subunits) as quantified by flow cytometry analysis of % intracellular ENaC/per cell (**Fig. 1A**, β subunit shown). ENaC protein expression was also higher in CF MDMs at baseline and persisted during infection with the CF intracellular pathogen *Burkholderia cenocepacia*, as compared to non-CF (**Fig. 1B**, γ subunit shown with corresponding densitometry, full blots in **Supplemental Fig 1**). Interestingly, ENaC levels decreased during infection in non-CF MDMs, which suggests a changing utilization pattern during infection in non-CF conditions. Confocal microscopy demonstrated that ENaC was widely expressed in the CF macrophage, with increased localization peripherally suggestive of localization to the plasma membrane. There was also inverse expression to CFTR, which was increased in non-CF and decreased in CF MDMs (**Fig. 1C**, ENaC puncta quantitation in **Fig. 1D**). Increased ENaC expression in CF was verified via direct CFTR pharmacologic inhibition (inh172) in non-CF MDMs. qRT-PCR of ENaC β subunit verified over-expression in CFTR inhibited cells, similar to primary human CF macrophages (homozygous F508del, **Fig. 1E**). However, ENaC was less abundantly expressed in CF MDMs compared to human CF airway epithelial cells (control homozygous F508del, **Fig. 1E**). We also verified that ENaC was present in other immune cells including granulocytes, and to a lesser extent in lymphocytes (**Fig. 1F**).

**Figure 1:**
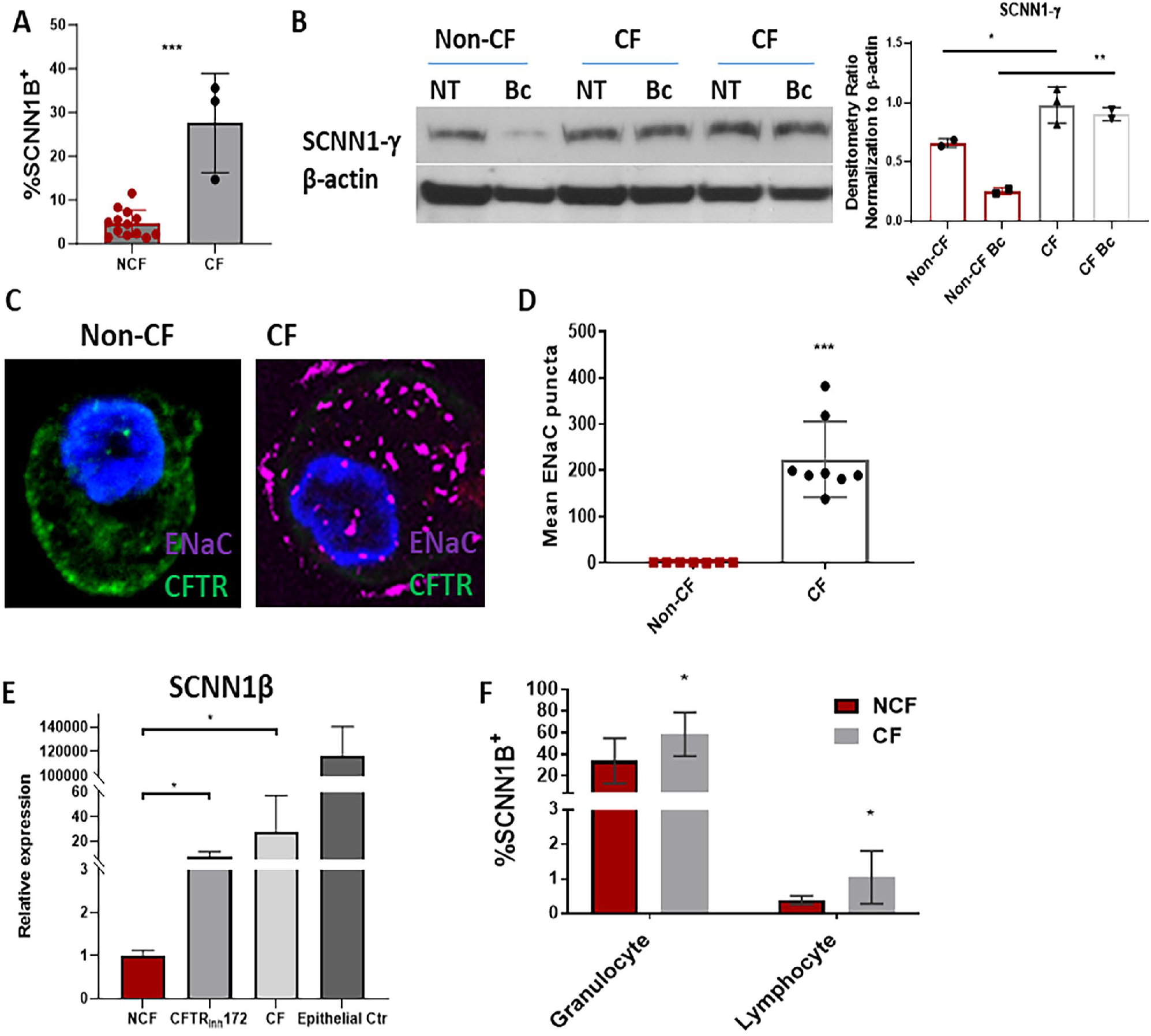
ENaC is over-expressed in CF immune cells. A) Flow cytometry intracellular detection of SCNN1β antibody in human non-CF and CF (F508del) MDMs. B) Western blot detection of SCNN1γ in human non-CF and CF MDMs at baseline (NT) and during infection with *B. cenocepacia* (Bc). Densitometry normalized to loading control. C) Confocal microscopy detection of ENaC (purple) and CFTR (green) basal expression in non-CF and CF MDMs. Nucleus stained blue with DAPI. Representative images shown. D) Quantitative mean ENaC puncta scoring for non-CF and CF MDMs from 1C, n=8. E) qRT-PCR SCNN1β expression in non-CF MDMs, non-CF MDMs with CFTR inhibition (CFTRinh172), CF MDMs, and compared to CF airway epithelial cells (F508del). F) SCNN1β expression in non-CF and CF granulocytes and lymphocytes via flow cytometry with statistical comparison within each cell type. “*” = p <0.05, “**” = p <0.01, “***” = p <0.001, via unpaired t-test for parametric data or Mann-Whitney U test for non-parametric data in 1A, 1D, 1F, n=4-8 individual donors for all experiments. One-way ANOVA used for 1B, 1E.

### Therapeutic manipulation influences ion channel expression in CF macrophages

Because of ENaC’s known associations with CFTR, we next investigated therapeutic manipulation of CFTR and its influence on macrophage ENaC expression. Recently, the highly effective CFTR modulator therapy ETI was approved for pwCF. Treatment of CF MDMs with ETI ex vivo reduced total ENaC abundance to near non-CF levels (**Figs. 2A**, β & Δ subunits).

**Figure 2:**
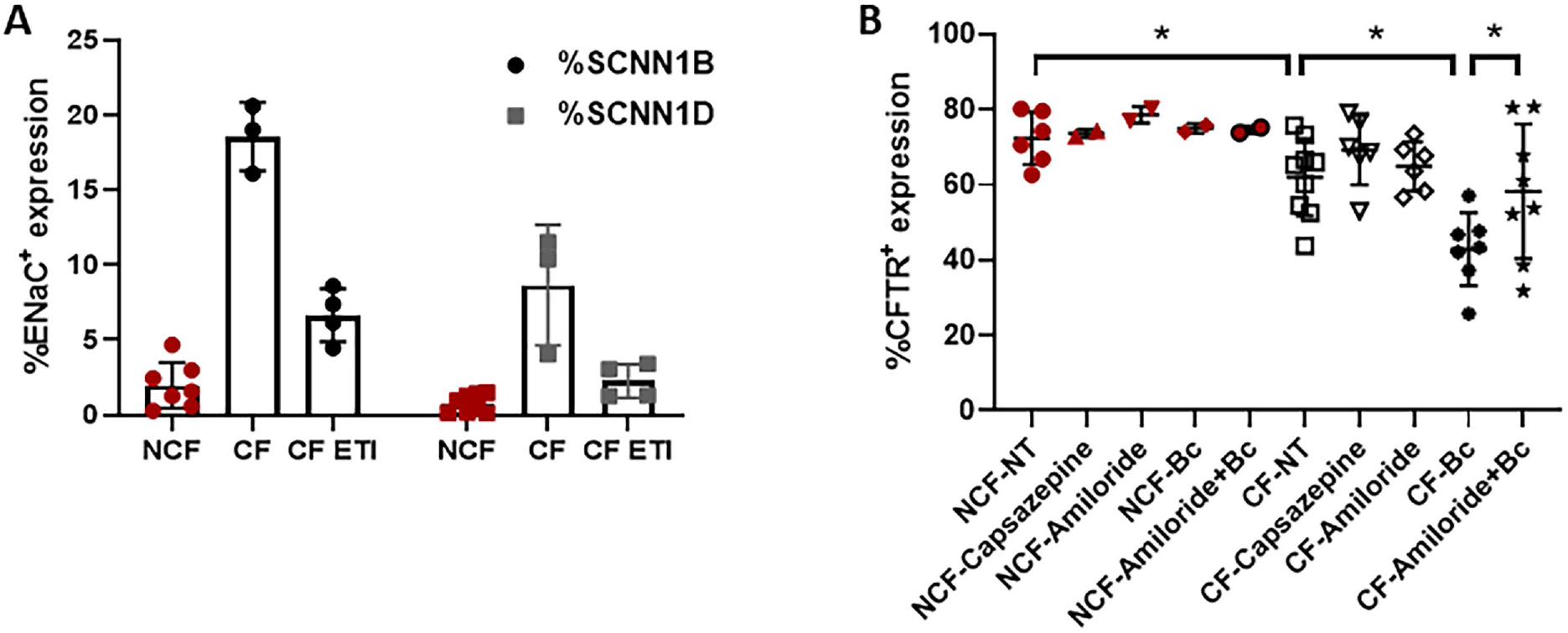
ENaC interactions with CFTR. A) Flow cytometry intracellular detection of SCNN1β or SCNN1Δ antibody in human non-CF and CF (F508del) MDMs alone or treated with ETI, n=3-7. B) Flow cytometry intracellular detection of CFTR in non-CF and CF MDMs in response to treatment with amiloride, capsazepine, infection with *B. cenocepacia*, or combinations, n=3-7, one-way ANOVA. “*” = p <0.05.

Conversely, ENaC inhibition alone with the pharmacologic inhibitor amiloride or ENaC activation with capsazepine did not alter CF or non-CF MDM CFTR (**Fig. 2B**). However, ENaC inhibition with amiloride did increase CF MDM CFTR expression during infection with *B. cenocepacia* compared to infected but untreated CF MDMs, (**Fig. 2B**, amiloride + Bc), suggesting activation during infectious stimulation.

There are many ion channels present in macrophages in addition to CFTR and ENaC,^18^ and amiloride can non-specifically impact channels besides ENaC. However, a targeted approach to ion channel manipulation is likely necessary for successful therapy development in CF. To gain an understanding of how ion channels are expressed in CF macrophages and how they respond to ENaC inhibition, we used a custom 48 gene Taqman microarray of ion channel expression during several conditions. Baseline expression levels for ion channels in CF MDMs relative to non-CF MDMs are shown in **Figure 3A**, grouped by ions regulated (chloride, sodium, calcium, potassium, or multiple). CF MDMs had higher expression of sodium channels SCNN1A (ENaC) and SCN7A, calcium channel CACNB1, and potassium channels KCNA3 and KCNE1 compared to non-CF MDMs (**Fig. 3A**).

**Figure 3:**
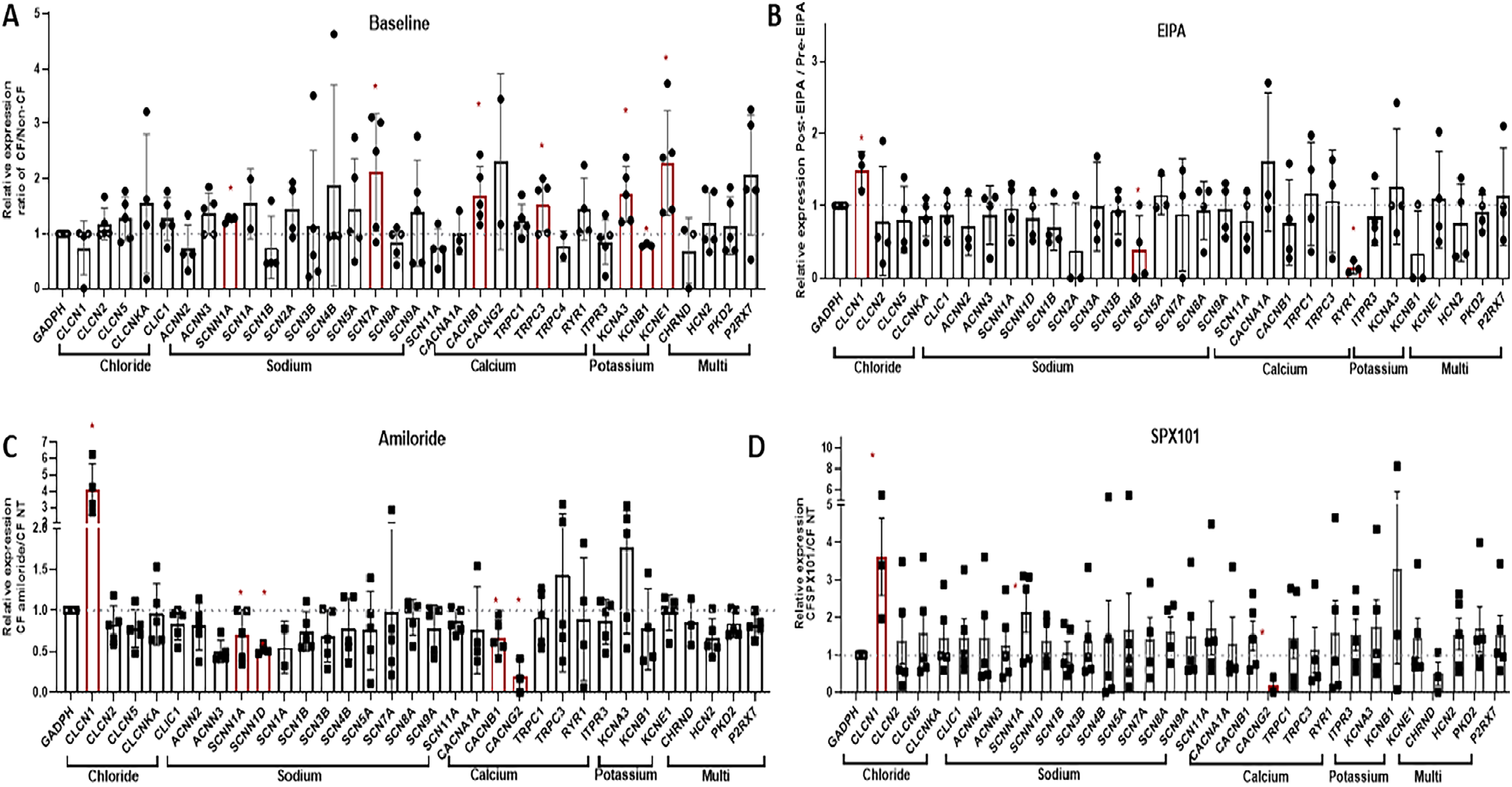
Compensatory ion channel changes. Custom taqman ion channel array for changes in non-CFTR ion channels at A) baseline, B) in response to EIPA broad sodium channel inhibition, C) Amiloride ENaC inhibition, and D) treatment with ENaC inhibitor SPX101. Channels are group by ion type. N=3-6. “*” = p <0.05, one-way ANOVA.

As a comparative control, we also treated CF MDMs with EIPA (broad sodium channel inhibitor) and found low expression of most sodium channels, but also compensatory increased expression of the chloride channel CLCN1 and the calcium channel RYR1 (**Fig. 3B**). Next, we treated CF MDMs with amiloride. After amiloride treatment, CF MDMs had lower expression of SCNN1A and SCNN1D (ENaC) and resolution of the previously shown altered expression of ion channels in **Fig. 3A** compared to untreated CF MDMs (**Fig. 3C**). Additionally, after amiloride treatment CF MDMs had increased CLCN1 and decreased CACNG2 compared to untreated CF MDMs (**Fig. 3C**). Results were similar when CF MDMs treated with amiloride were compared to treated non-CF MDMs, except for no change in KCNA3 (data not shown).

Further, ACNN, SCNN, and SCN channel expression were similar between CF and non-CF (data not shown). We then compared amiloride findings to that of SPX-101, a peptide mimetic of the natural regulator of ENaC activity by short palate, lung, and nasal epithelial clone 1 (SPLUNC1).^19^ Similar to amiloride, CF MDMs treated with SPX-101 demonstrated increased expression of CLCN1 and decreased expression of CACNG2 with normalization of ENaC expression (**Fig. 3D**). Overall, these data demonstrate baseline differences in CF ion channel expression, with minimal off-target effects during ENaC inhibition as compared to broad sodium channel inhibition.

### Functional consequences of ENaC inhibition in CF macrophages

We previously demonstrated that human CF macrophages are deficient in intracellular bacterial killing through failed NOX assembly and reduced autophagy.^20-24^ To investigate if ENaC contributes to CF macrophage NOX and autophagy dysfunction we performed studies of reactive oxygen species (ROS) production (as a surrogate of NOX assembly) and autophagy. Human CF (F508del, no CFTR modulator) and non-CF MDMs were infected with *B. cenocepacia* to induce an oxidative burst. ROS production was measured through a DCF assay. Similar to our prior work, we found that ROS production in CF MDMs was reduced 40% compared to non-CF (**Fig. 4A**, normalized to non-CF). When CF MDMs were treated with capsazepine, there was no significant change in ROS (**Fig. 4A**). In contrast, treatment with amiloride normalized ROS in CF MDMs compared to untreated CF (**Fig. 4A**), while ETI or SPX-101 treatment led to non-significant increases in ROS production compared to baseline. We further examined expression of cytosolic and membrane markers of NOX complex assembly at early (30 minutes) and late (4h) timepoints in response to the prior conditions. At 30 minutes, we found decreased membrane total and phosphorylated p47^*phox*^ during *B. cenocepacia* infection in CF, with slight changes in phosphorylated p47^*phox*^ in response to treatment with amiloride (densitometry in **Supplemental Fig. 2**, full blots **Supplemental Figs. 4-9**). Other NOX protein markers were unaffected by treatments. At 4 hours, membrane bound Rac2 and cytoplasmic phosphorylated p47^*phox*^ were increased in CF in response to PMA and amiloride or SPX-101, but other proteins remained unaffected (densitometry in **Supplemental Fig. 3**, full blots **Supplemental Figs. 4-9**).

We then examined macrophage autophagy using western blot to detect LC3-I to LC3-II conversion (autophagosome formation). As previously published, we saw a decrease in autophagy in response to infection in CF MDMs compared to non-CF (densitometry in **Fig. 4B**, blots in **Supplemental Fig 1**). However, the addition of amiloride to inhibit ENaC increased autophagy during infection in CF (**Fig. 4B**). We verified these findings with a flow cytometry-based autophagy assay, which demonstrated similar results (**Fig. 4C**). We also compared treatment with SPX-101 and ETI, which increased autophagy to similar levels compared to amiloride (**Fig. 4C**). Combinations of amiloride or SPX-101 did not show any further additive benefit to compounds alone (**Fig. 4C**).

**Figure 4:**
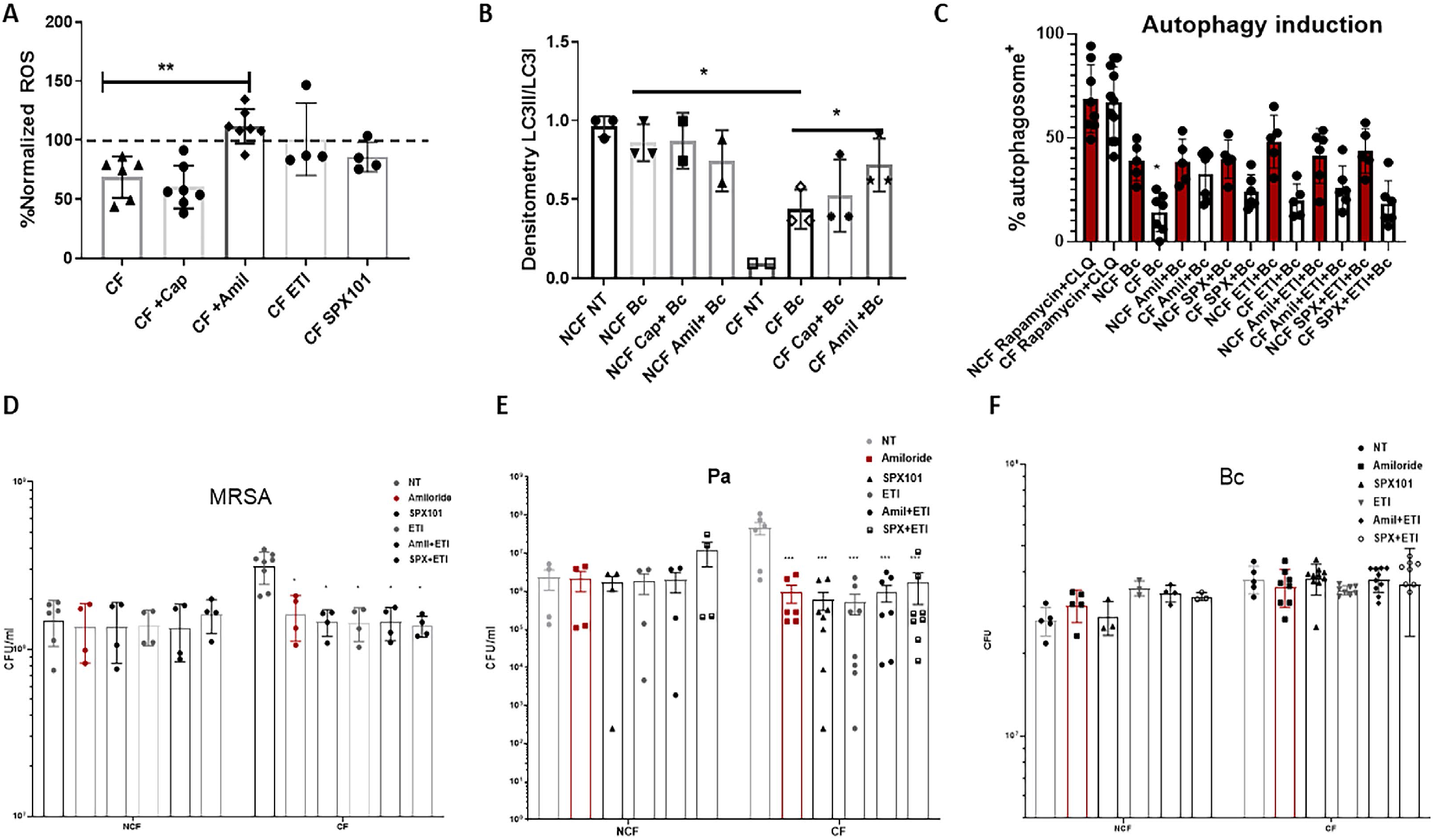
Phagocyte functional changes with ENaC inhibition. A) Changes in reactive oxygen species (ROS) production in CF MDMs normalized to non-CF (dashed line represents average non-CF response) in response to ENaC activators or inhibitors. B) Changes in autophagy induction in CF and non-CF MDMs during infection with *B. cenocepacia* +/-ENaC manipulation. Autophagy measured by LC3 II/I ratios via western blot. Summed densitometry shown. C) Changes in autophagy induction during infection with *B. cenocepacia* +/-ENaC manipulation. Autophagy measured by flow cytometry quantitative CytoID assay as confirmation of 4B. Changes in bacterial counts for D) MRSA, E) *Pseudomonas aeruginosa*, and F) *B. cenocepacia* in CF and non-CF MDMs in response to ENaC modulation or ETI. N=3-8, one-way ANOVA. “*” = p <0.05, “**” = p <0.01, “***” = p <0.001.

Next, we examined if the observed alterations in ROS production and autophagy induction altered bacterial killing. Interestingly, MRSA and *P. aeruginosa* bacterial load in CF MDMs were equally reduced by amiloride, SPX-101, ETI treatment, or combinations (**Fig. 4D-E**). Furthermore, there was no reduction in *B. cenocepacia* bacterial load with any treatment (**Fig. 4F**). Bacterial load in non-CF MDMs was unaffected by any treatments (**Fig. 4D-F**).

Last, we examined the impact of ENaC modulation upon CF macrophage cytokine production during persistent infection with *B. cenocepacia* and treatment with ETI, amiloride, SPX-101, or combinations. Prior to infection, MDMs were exposed to ASN to mimic the CF airway milieu, which can impact cytokine secretion and phagocytosis.^5,25^ Overall, we found no significant difference between treatments, as all treatments decreased IL-1β, IL-10, and IL-23 production by CF macrophages (**Fig. 5**). There were minor variations between treatments for reductions in TNF-α, IL-12p70, and increased MCP-1. IFN-γ showed a trend towards reduced levels for all treatments. IFN-a2, IL-6, IL-8, IL-17a, IL-18, and IL-33 were unchanged by any treatment (**Fig. 5)**.

**Figure 5:**
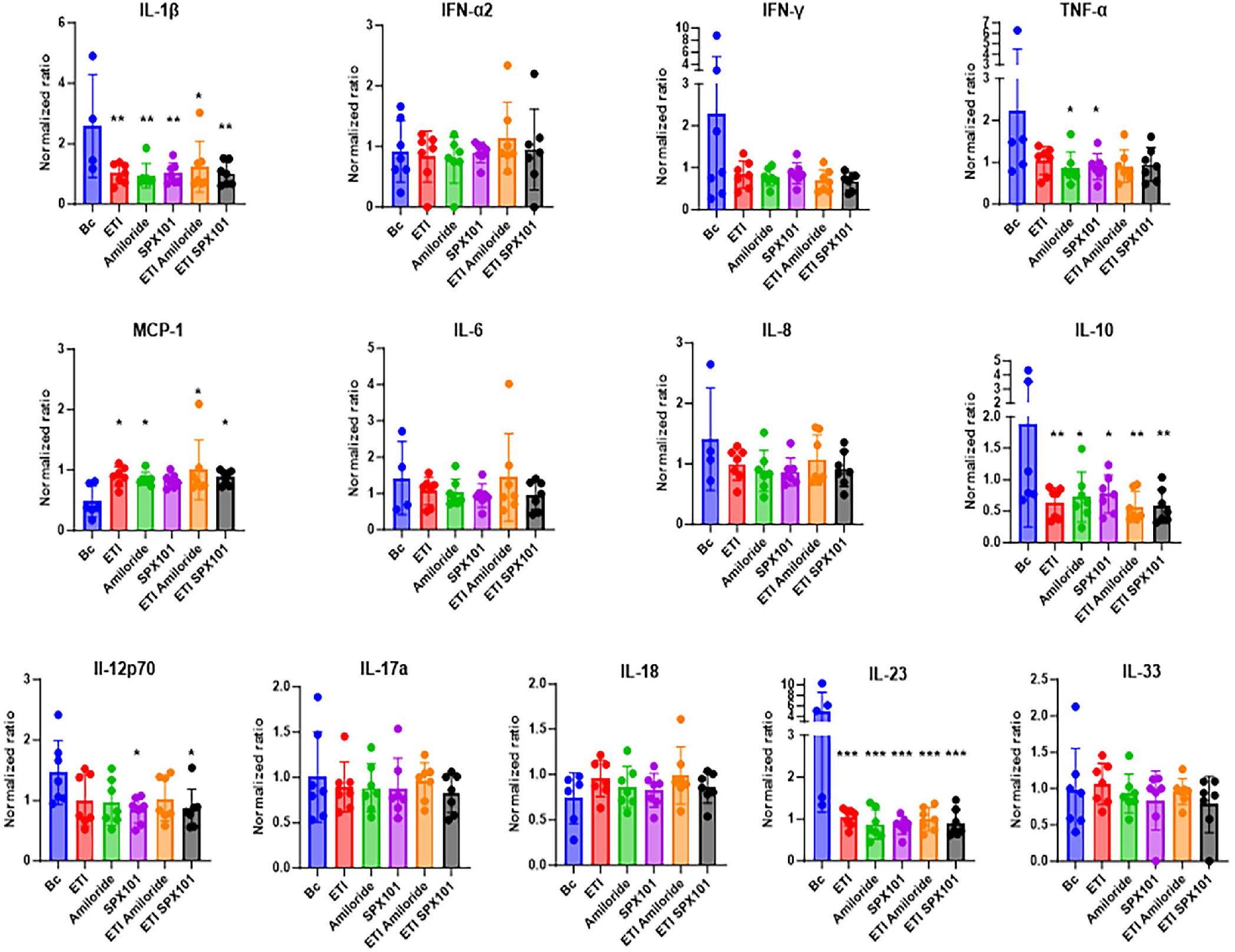
Cytokine responses to ENaC manipulation. Multiplex cytokine assay for MDMs during B. cenocepacia infection or treatment with ETI, amiloride, SPX-101, or combinations. CF results shown normalized to non-CF. n=5-7, one-way ANOVA. “*” = p <0.05, “**” = p <0.01, “***” = p <0.001.

## DISCUSSION

The role of ion channel dysfunction in relation to dysregulated immunity in pwCF remains unclear. Further, the interplay between ENaC and CFTR is a complex landscape based on interconnected signaling. Our findings describe an overall role for ENaC in concert with CFTR in the regulation of macrophage function (**Fig 6**). We found that ENaC was broadly expressed in a variety of immune cells and specifically overexpressed in CF immune cells. As shown in Figure 6, ENaC has a direct effect on several known areas of macrophage dysfunction in CF, including ROS production, autophagy induction, and cytokine secretion. Whether ENaC manipulation can be a CFTR agnostic therapy with beneficial effects on chronic inflammation and infection needs further study but shows promise for non-CFTR ion-channel mediated effects in immune cells. Ongoing airway based ENaC inhibition studies in CF continue to show limited benefit^26^, but future trials may need to be targeted towards immune cells.

**Figure 6:**
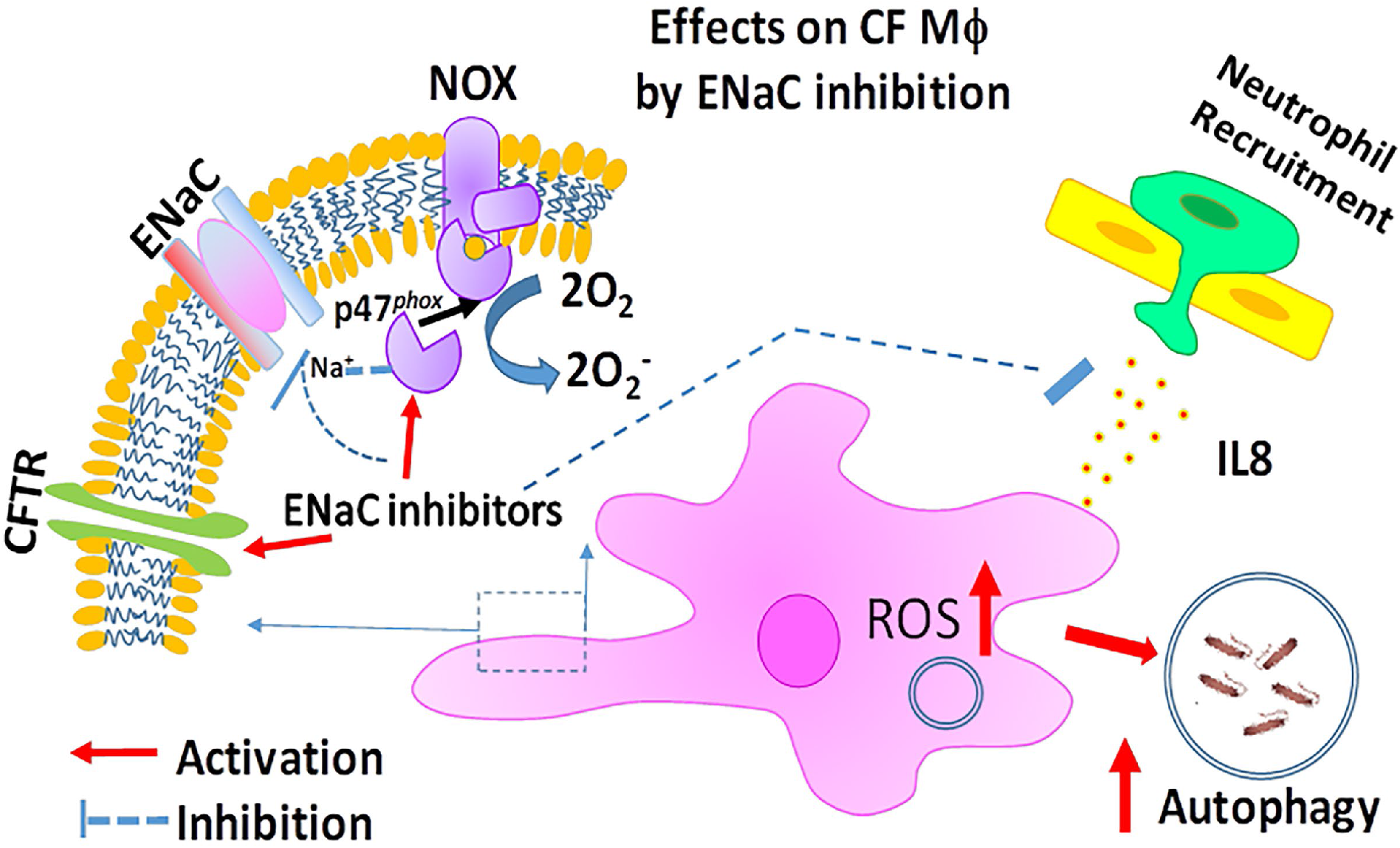
Graphical summary of results.

ENaC over-expression in CF immune cells presents a new aspect of CF pathology, particularly concerning immune responses. Our findings revealed that ENaC is markedly upregulated in CF macrophages compared to non-CF controls, as demonstrated through various methods including western blot, flow cytometry, and qRT-PCR. This heightened expression persists during infection with the CF pathogen *B. cenocepacia*, suggesting a potential role of ENaC in exacerbating persistent infection. Notably, while ENaC levels decrease in non-CF macrophages during infection, indicating a modifiable pattern in response to pathogen exposure, CF macrophages maintain elevated ENaC levels, potentially contributing to persistent immune challenges. The observed inverse relationship between ENaC and CFTR in CF macrophages— where CFTR is diminished and ENaC is increased—further underscores the interplay between these channels. Pharmacologic inhibition of CFTR, which results in increased ENaC expression in non-CF macrophages, mirrors the over-expression seen in CF MDMs, reinforcing the idea of a regulatory interaction between CFTR and ENaC. Treatment with ETI reduced ENaC levels in CF macrophages, aligning them closer to non-CF levels. In contrast, selective ENaC inhibition with amiloride or activation with capsazepine did not significantly impact CFTR expression, though amiloride did enhance CFTR levels during infection, suggesting a dynamic response to ion channel modulation. How long-term use of CFTR modulators or other ion channel regulators will influence ENaC in CF immune cells remains to be determined. Additionally, continued mechanistic studies of cellular sodium signaling in CF macrophages are warranted. Further, our findings highlight the necessity of targeted ion channel therapies to avoid broad off-target effects and optimize treatment efficacy.

Recently, ENaC mediated sodium influx was also found to exacerbate NLRP3-dependent inflammation in CF monocytes and epithelia as evidenced by increased levels of IL-18, IL-1β, and caspase-1 activity.^12^ The authors found that pre-treatment with NLRP3 inflammasome inhibitors could reverse these defects. Functionally, we found that ENaC inhibition with amiloride or the use of the ENaC modulator SPX-101 resulted in normalized ROS production and increased autophagy in CF macrophages, addressing previously observed deficiencies in NOX assembly and autophagy. Despite these improvements in immune function, bacterial clearance remained unaltered for certain pathogens, which suggests that while ENaC modulation can ameliorate some aspects of macrophage dysfunction, it may not fully restore effective pathogen clearance. Viral infection was not examined in this study, but recent evidence demonstrates that the SARS-CoV-2 spike protein activates ENaC through proteolytic cleavage of side chains.^27^ How acute viral infections in the setting of chronic ENaC hyperactivation in CF alters immune response is another target for further study.

Cytokine production by CF macrophages was generally unaffected by the treatments studied herein, indicating that while ion channel modulation can influence some immune parameters, it may not comprehensively address all aspects of CF immune dysregulation. Overall, these insights into ENaC and CFTR interactions not only advance our understanding of CF macrophage pathology but also inform future therapeutic strategies aiming to correct immune deficiencies in CF, which may need to involve combinatorial approaches.

## ACKNOWLEDGEMENTS

Thank you to Robert Tarran, PhD for donation of SPX-101 and Martina Gentzsch, PhD, University of North Carolina – Chapel Hill, and Cystic Fibrosis Foundation CFTR antibody distribution program for CFTR antibodies.

## DISCLOSURES

The authors have no relevant disclosures.

## Notes

**Grant support:** This study was supported by CF Foundation grant KOPP20I0 (BTK) and NIH R01 HL158747 (BTK, SPS) and R01 HL148171 (BTK, EB). Cure CF Columbus Immune Core grant support provided by The Ohio State University Center for Clinical and Translational Science (National Center for Advancing Translational Sciences, Grant UL1TR002733) and by the CF Foundation (Research Development Program, Grant MCCOY19RO). Human subject samples were additionally provided by the CF Biospecimen repository at the Children’s Healthcare of Atlanta and Emory University Discovery Core (P30-DK125013) and Immunophenotyping Core.

### Competing Interest Statement

The authors have declared no competing interest.

