## Supplementary figures and images for "ENaC contributes to macrophage dysfunction in cystic fibrosis"

### Supplemental figures

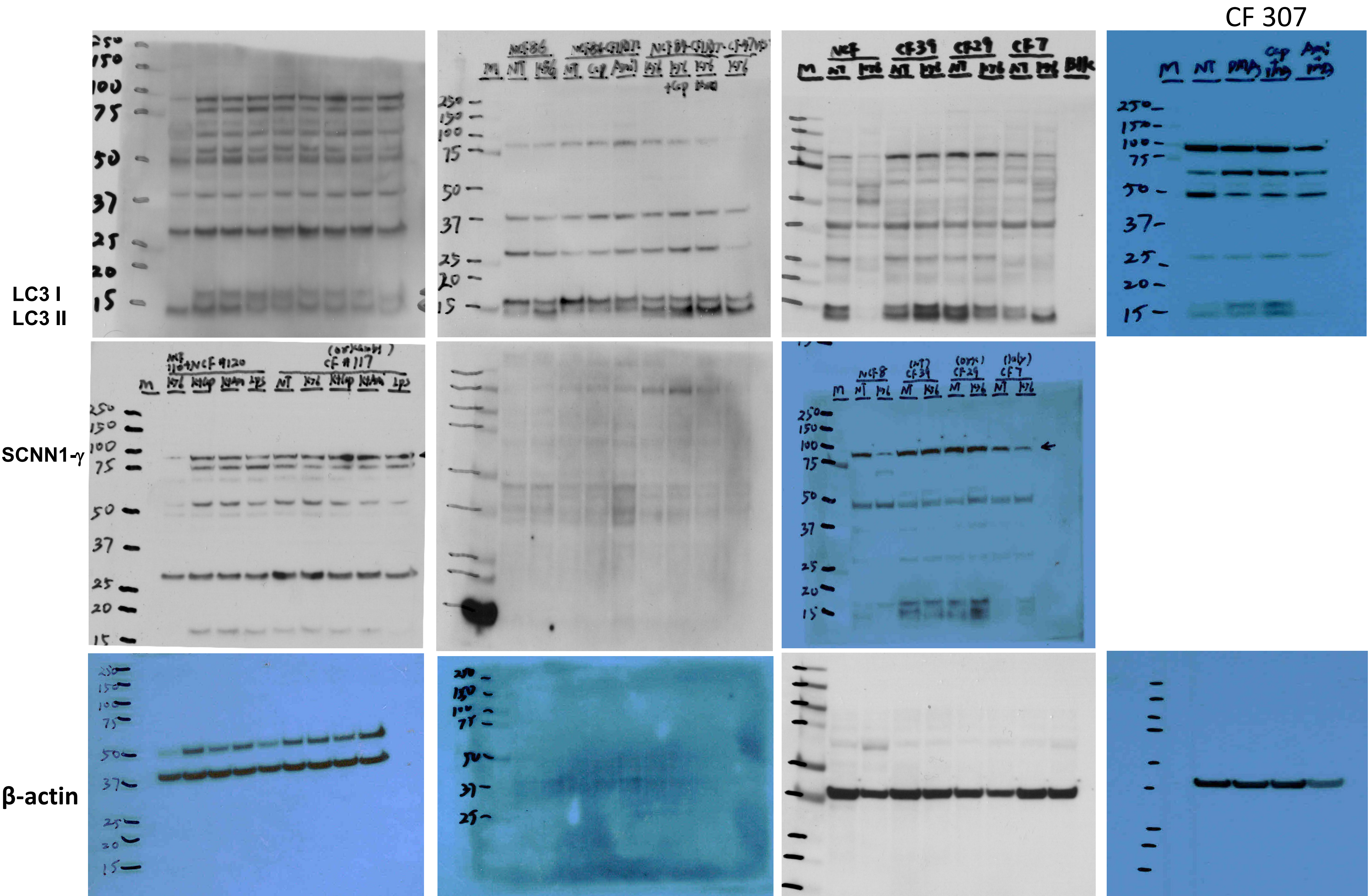

#### 30 minutes western densitometry

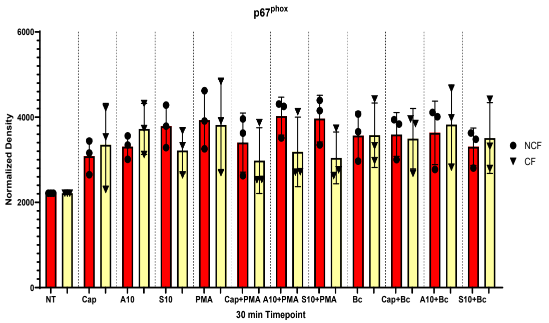


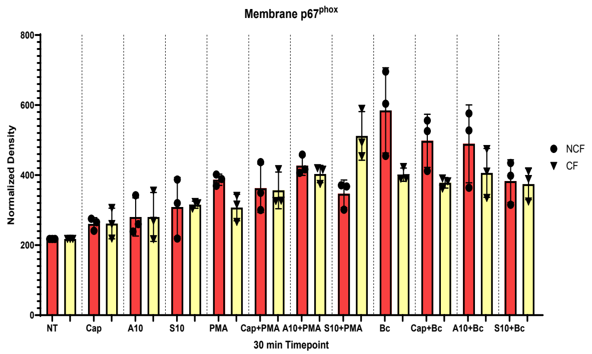


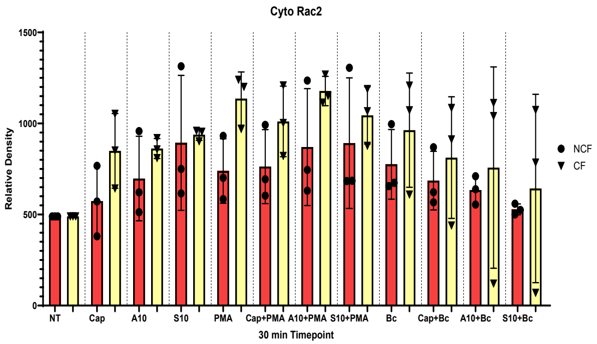


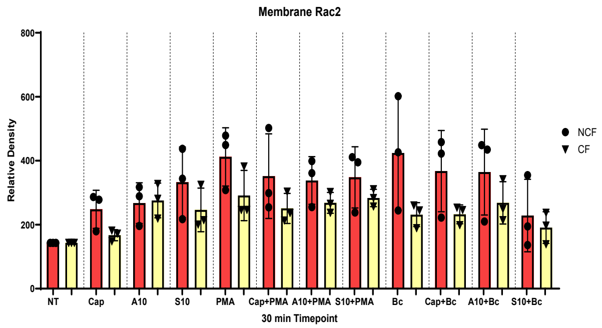


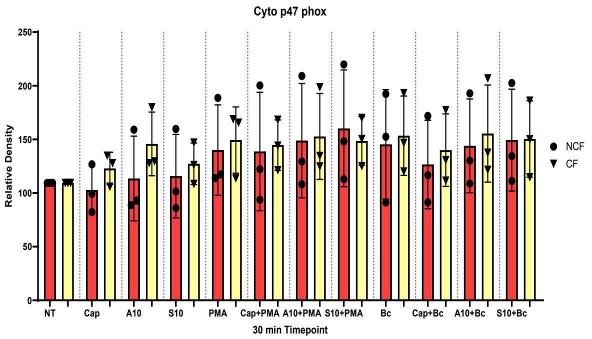


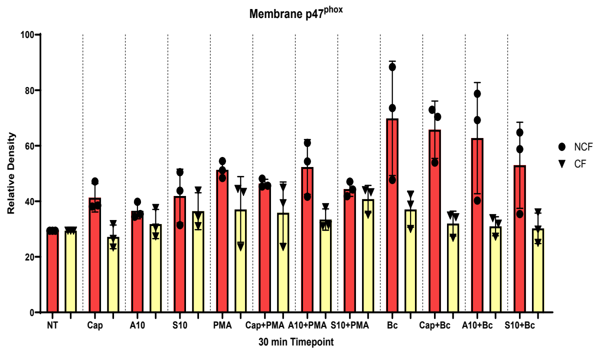


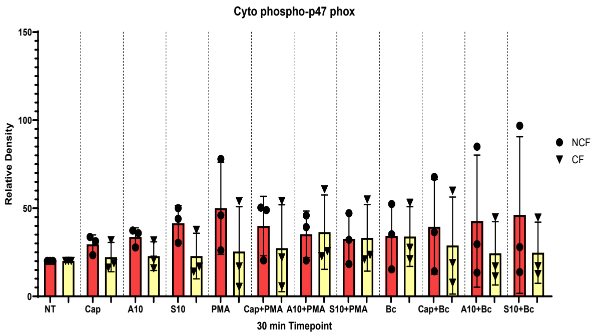


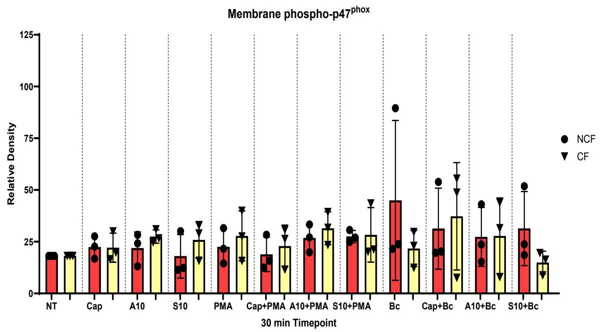


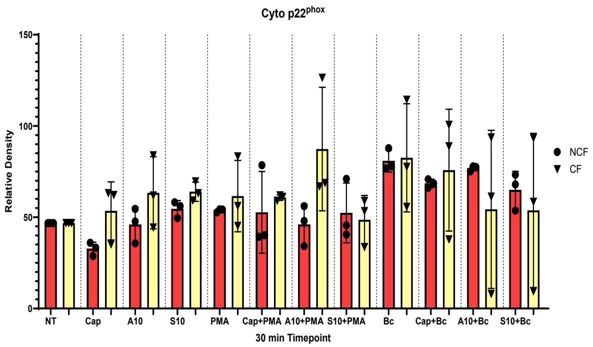


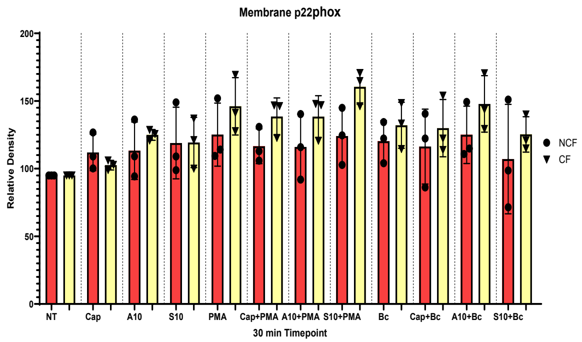

#### 4 hour western densitometry

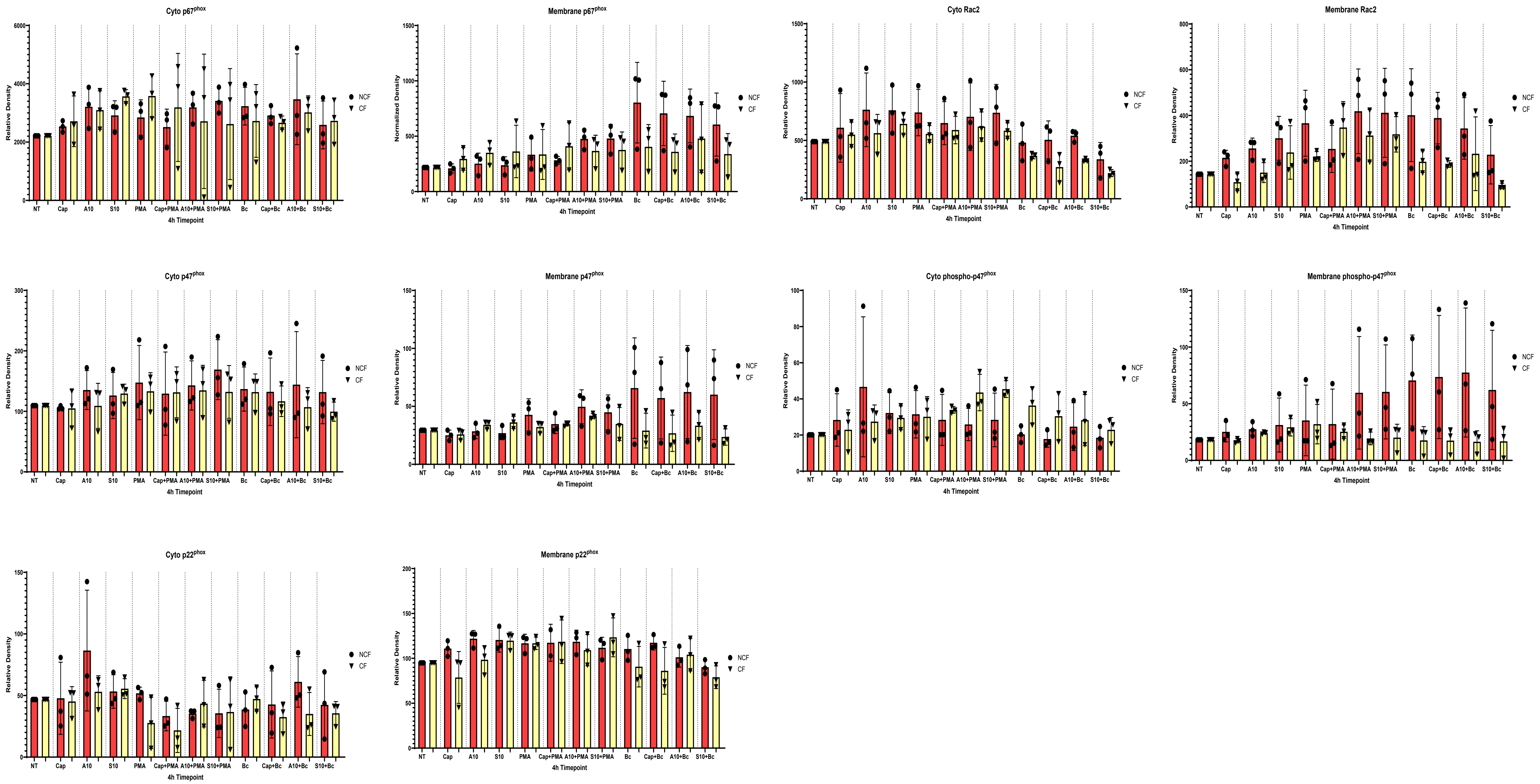

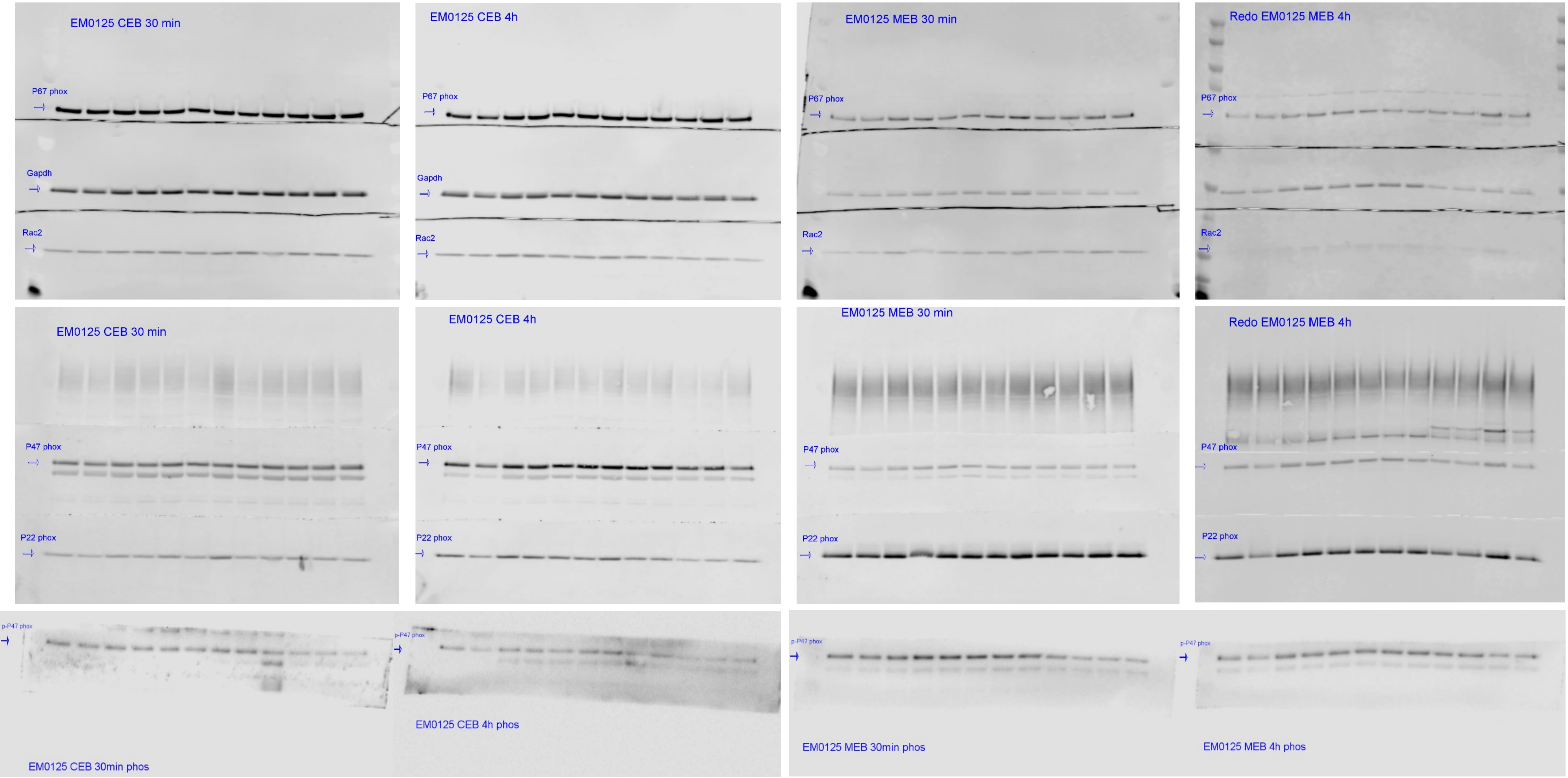

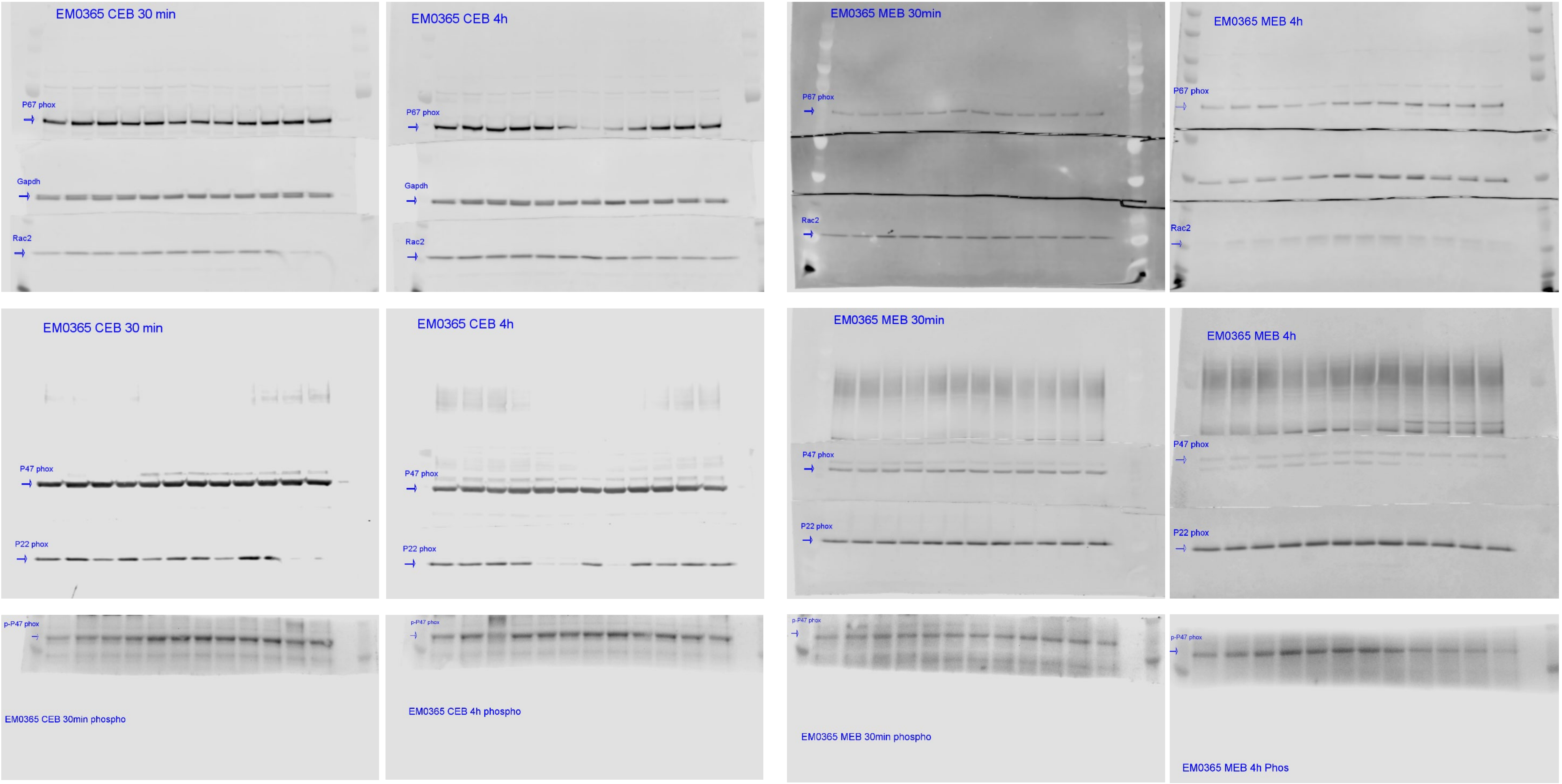

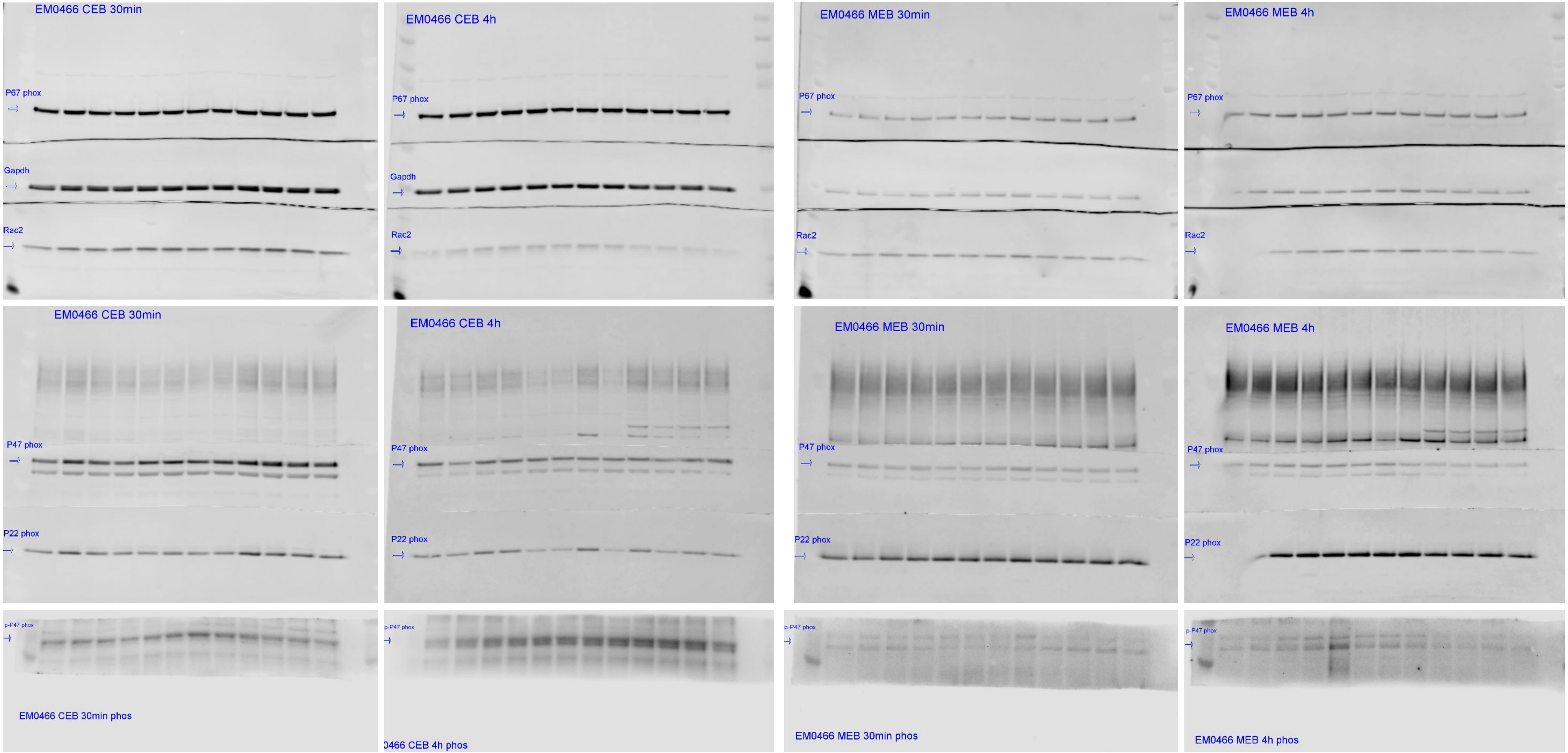

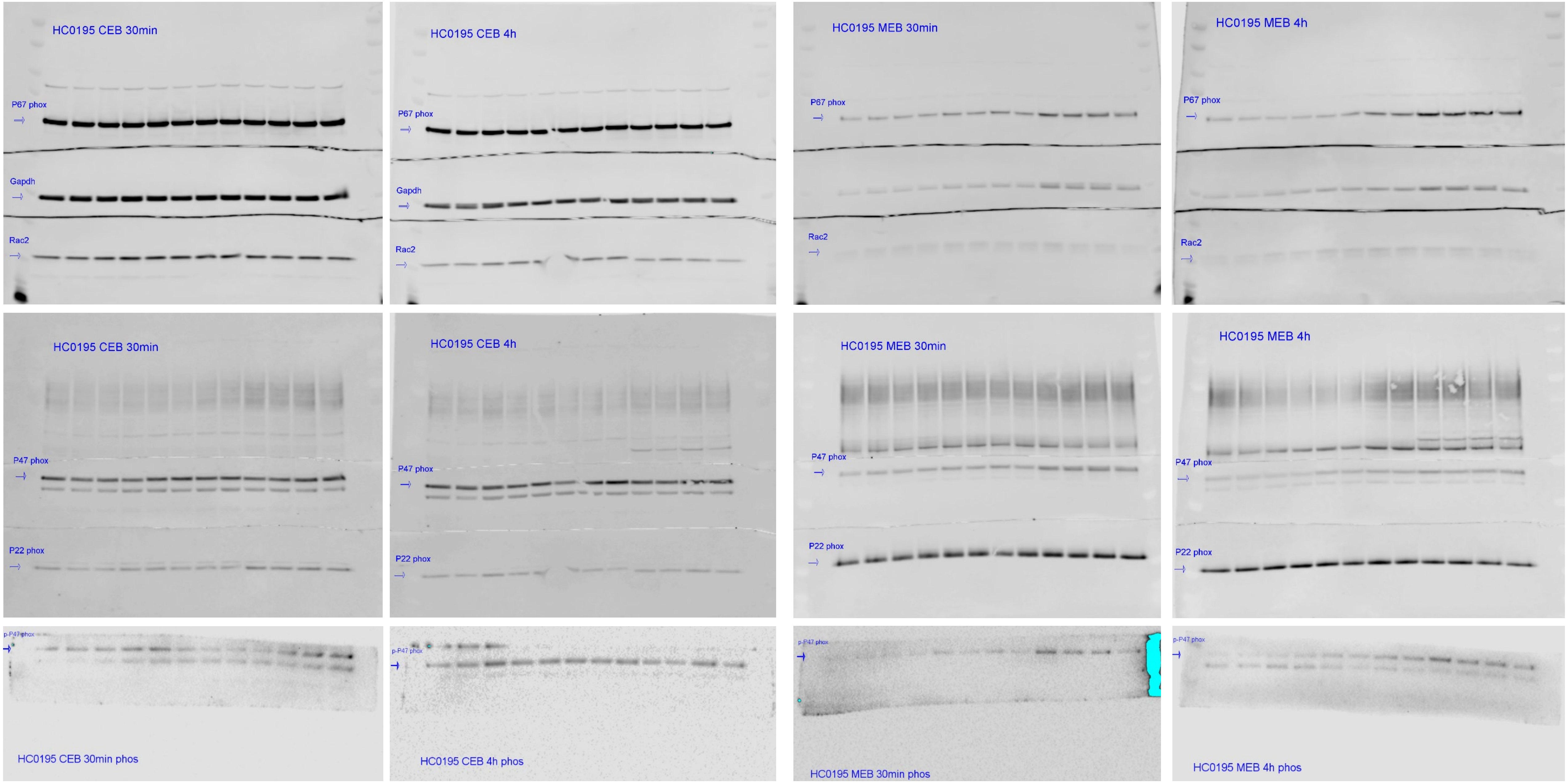

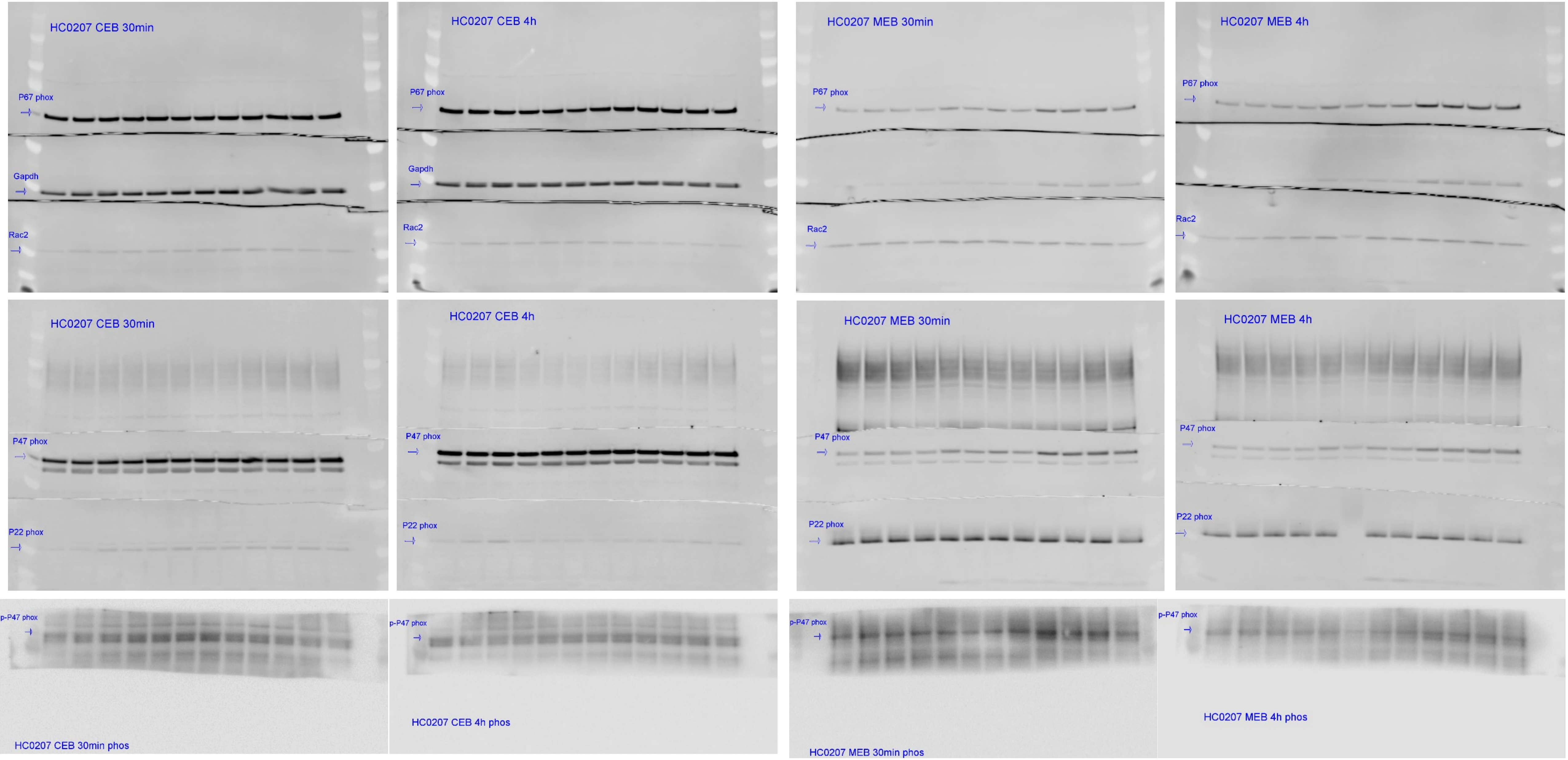

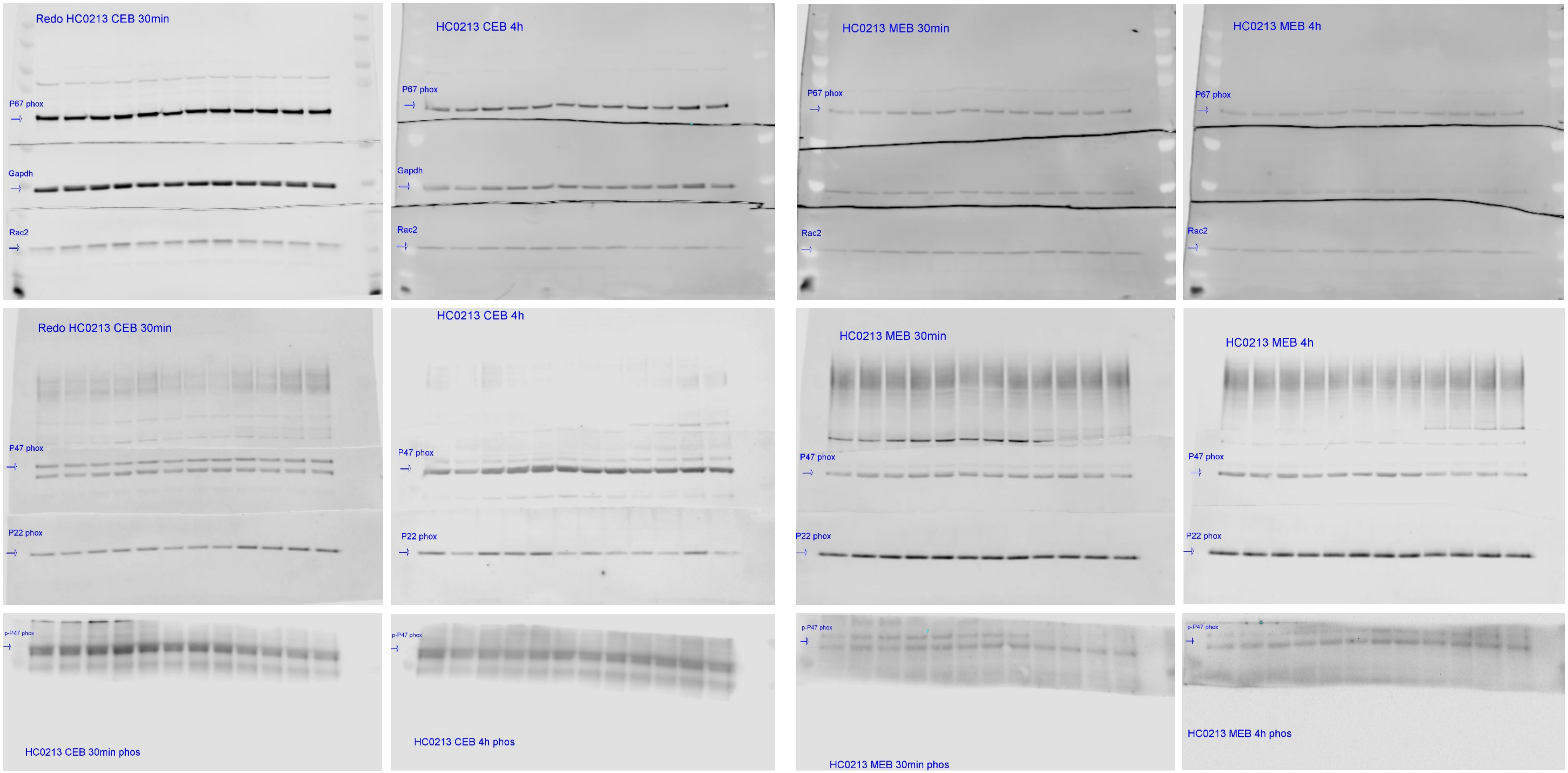
